## Supplemental figure for "Root-specific secondary metabolism at the single-cell level: a case study of theanine metabolism and regulation in the roots of tea plants (*Camellia sinensis*)"

### Supplemental Figures

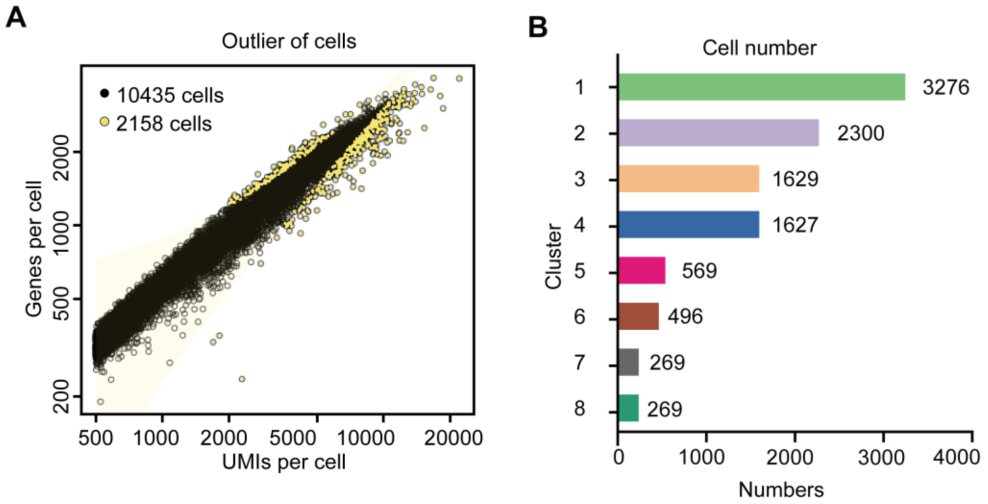


**Supplemental Figure 1. Summary the tea roots scRNA-seq. (A)** Filter delocalized cell maps fitted to generalized linear models. The horizontal axis is the unique molecular identifiers (UMI) number in each cell, and the vertical axis is the gene number in each cell. According to the linear relationship between them, the distribution model is fitted. The yellow points represent delocalized cells, which will be removed in the follow-up analysis. **(B)** The cell number in each cell clusters of tea roots scRNA-seq.

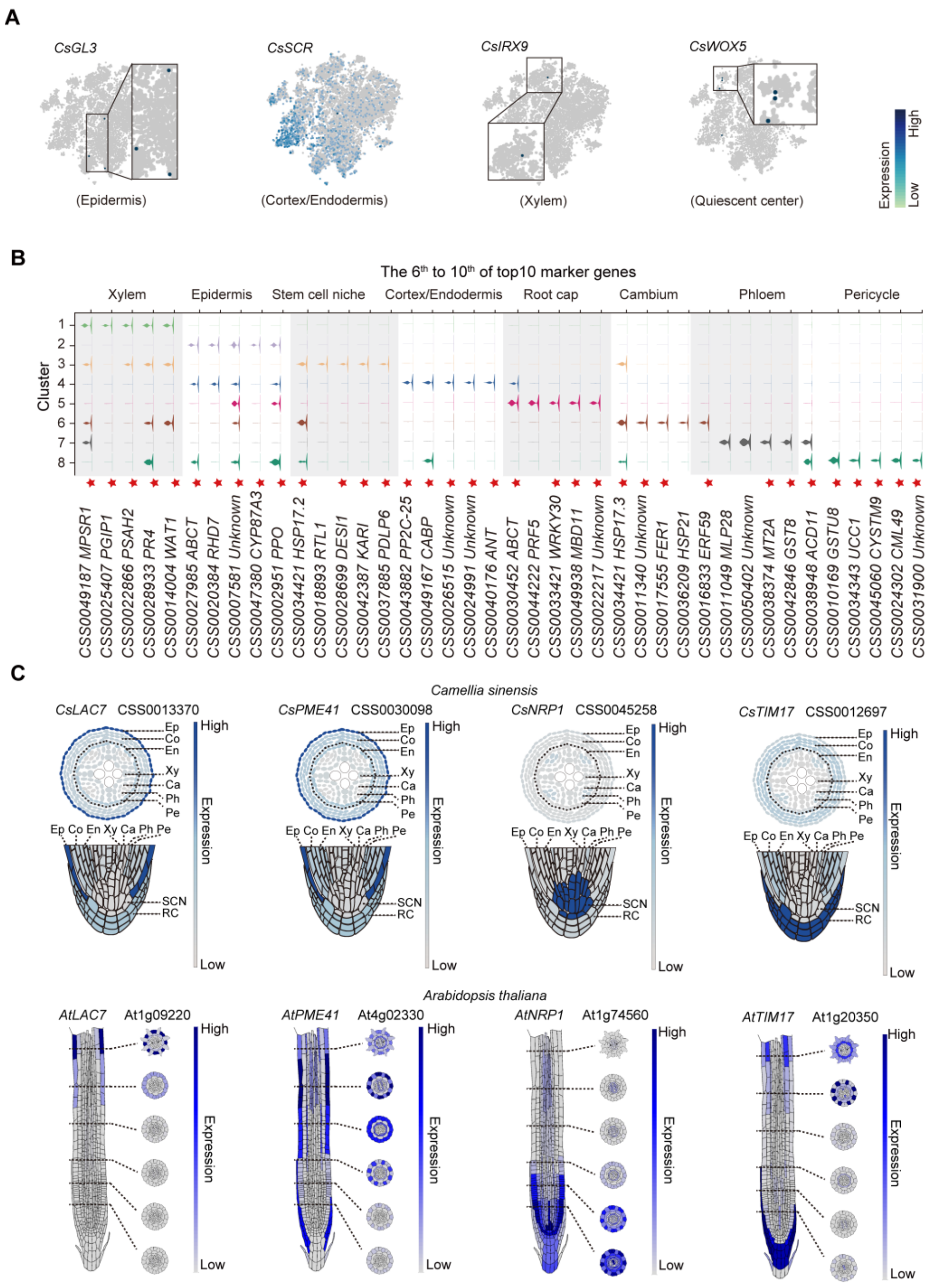


**Supplemental Figure 2. Cell cluster annotation. (A)** *t*-SNE visualization of cell-type marker genes (*CsGL3, CsSCR, CsIRX9 and CsWOX5*) of root cells. Color bar indicated gene expression level. **(B)** Violin plot showing expression patterns of 6$^{th}$ to 10$^{th}$ in the top10 differential marker genes of each cell cluster. The height and width of the violin represent the gene expression level and the proportion of cells expressing in the cluster, respectively. Asterisks represent homologous marker genes that are present in PlantscRNAdb. **(C)** The tissue-located heatmap of *LAC7, PME41, NRP1* and *TIM17*. The color bar illustrated gene expression level for the scRNA-seq data. The *Arabidopsis thaliana* data and pictures were generated from the Root Cell Atlas (https://rootcellatlas.org/).

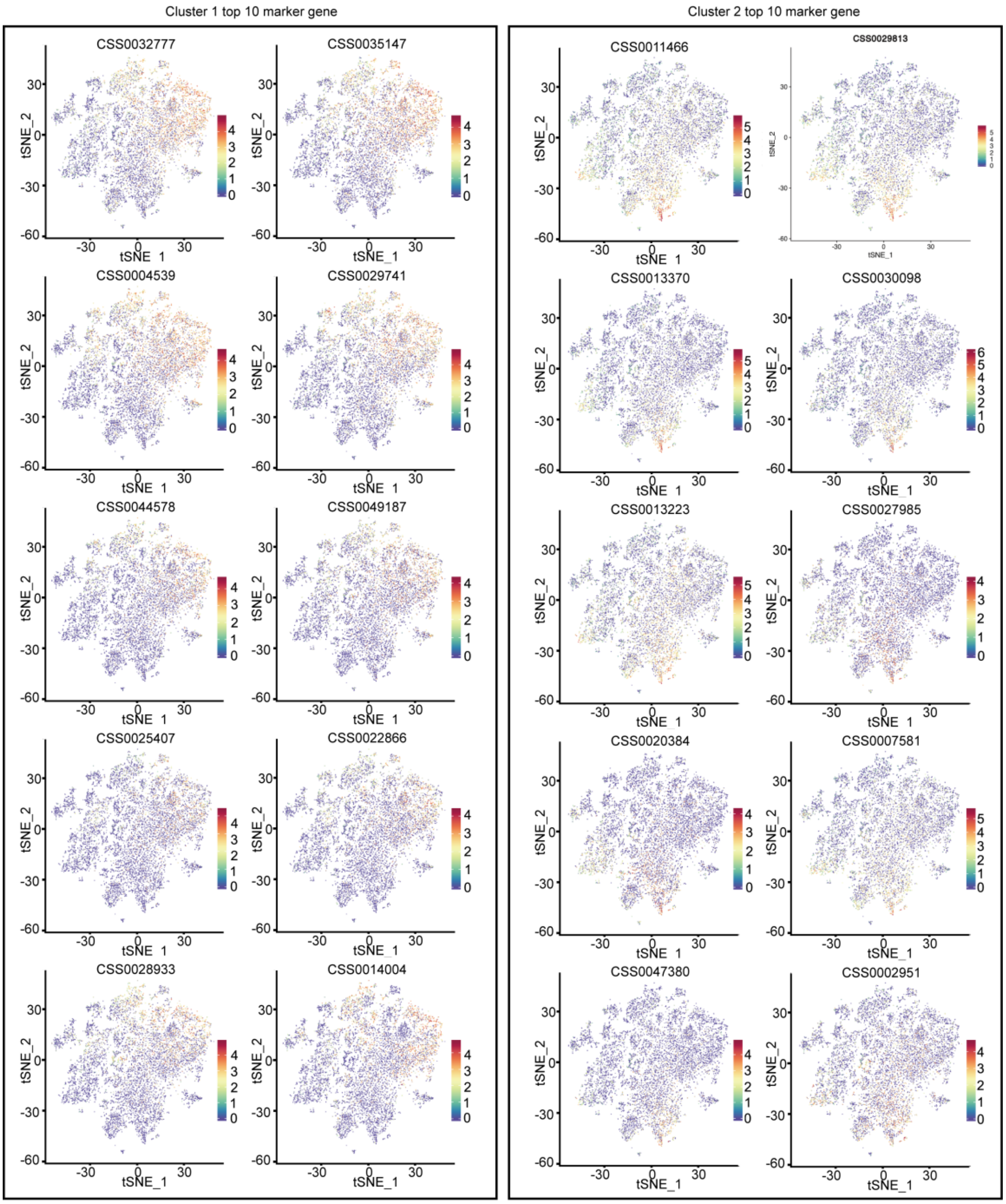


**Supplemental Figure 3. *t*-SNE visualization of cluster 1 and cluster 2 top 10 marker genes.** *t*-SNE plots show the transcript accumulation of cluster 1 and cluster 2 top 10 marker genes in individual cells. Color intensity indicates the relative transcript level for the indicated gene in each cell.

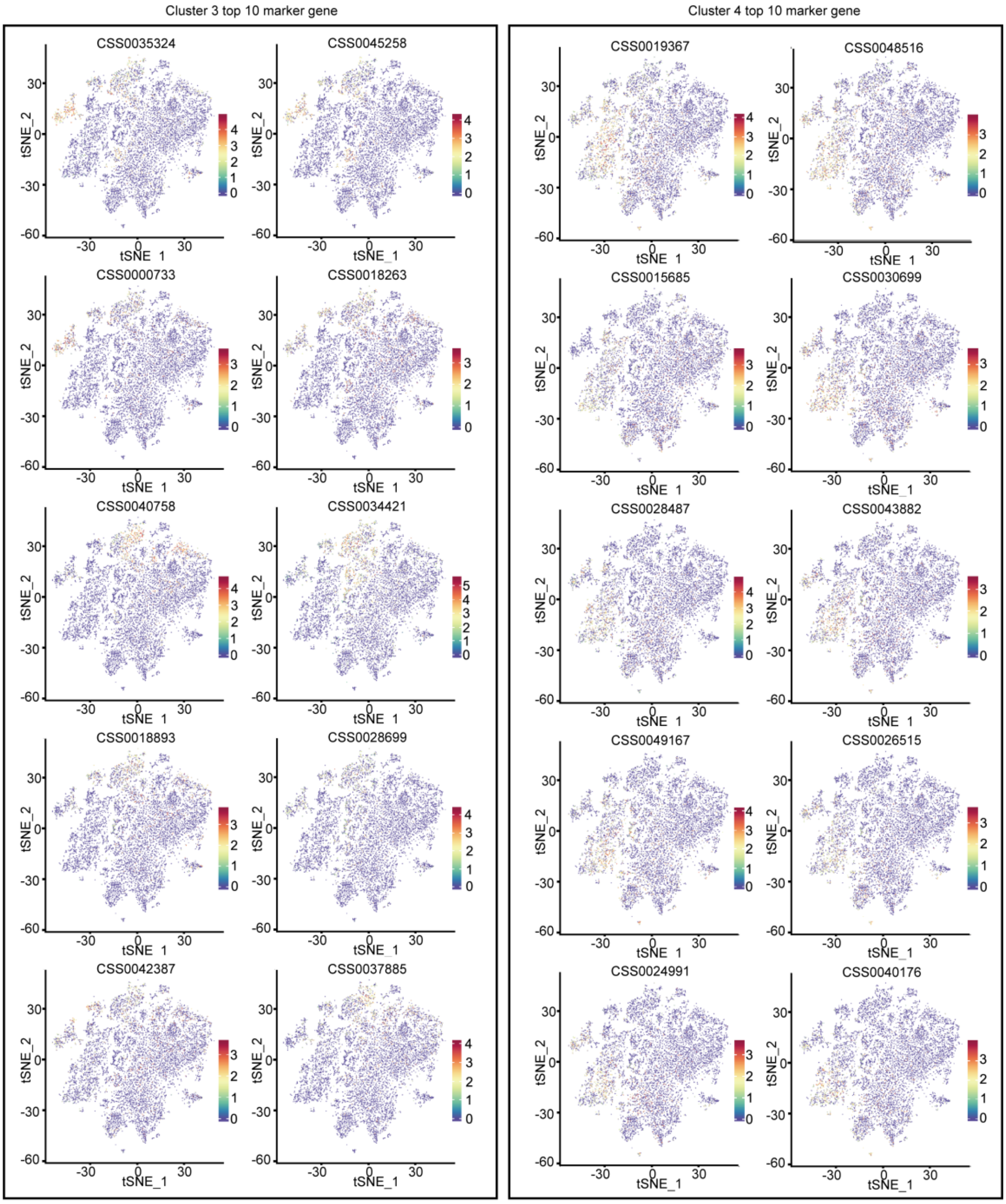


**Supplemental Figure 4. *t*-SNE visualization of cluster 3 and cluster 4 top 10 marker genes.** *t*-SNE plots shows the transcript accumulation of cluster 3 and cluster 4 top 10 marker genes in individual cells. Color intensity indicates the relative transcript level for the indicated gene in each cell.

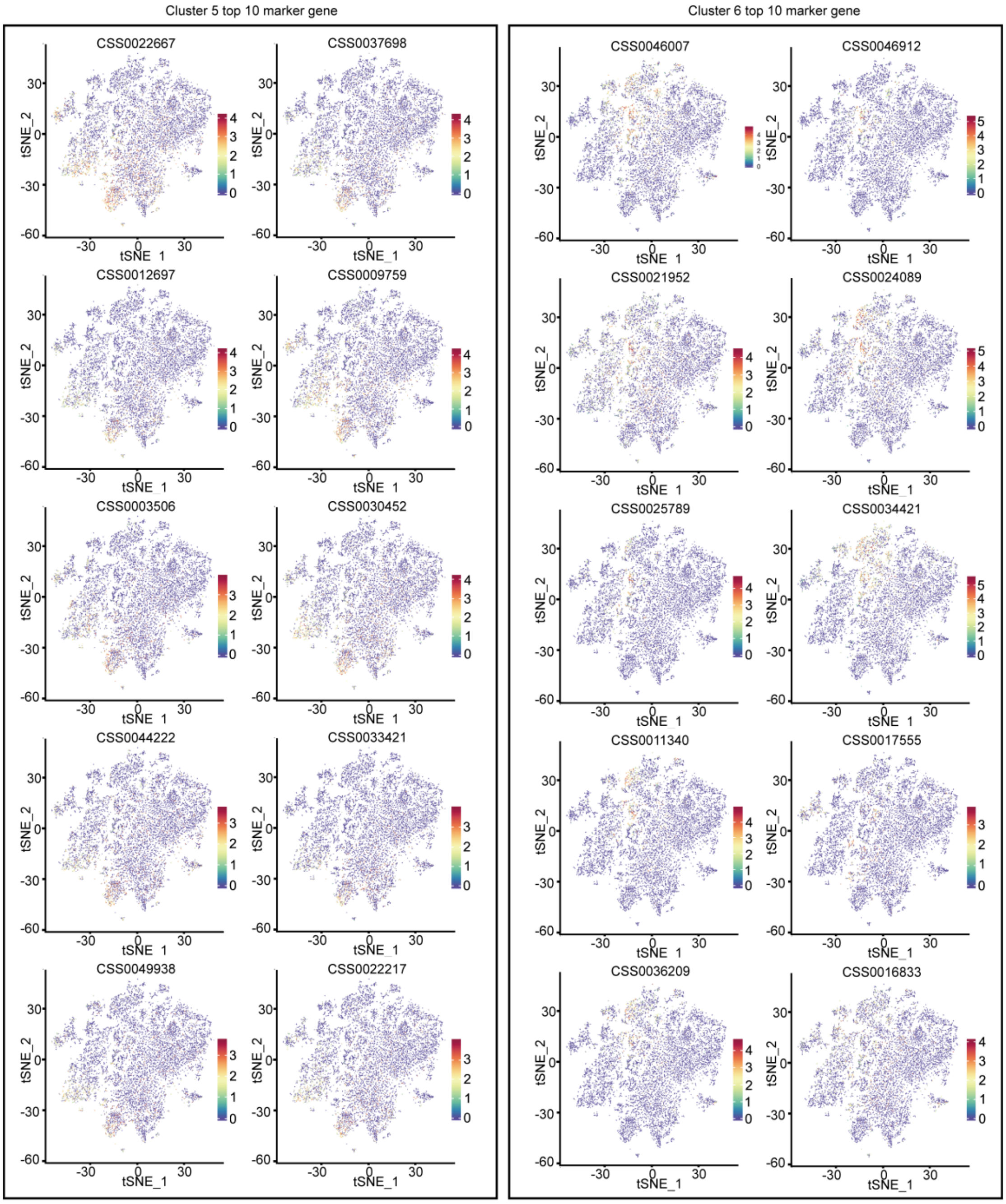


**Supplemental Figure 5. *t*-SNE visualization of cluster 5 and cluster 6 top 10 marker genes.** *t*-SNE plots shows the transcript accumulation of cluster 5 and cluster 6 top 10 marker genes in individual cells. Color intensity indicates the relative transcript level for the indicated gene in each cell.

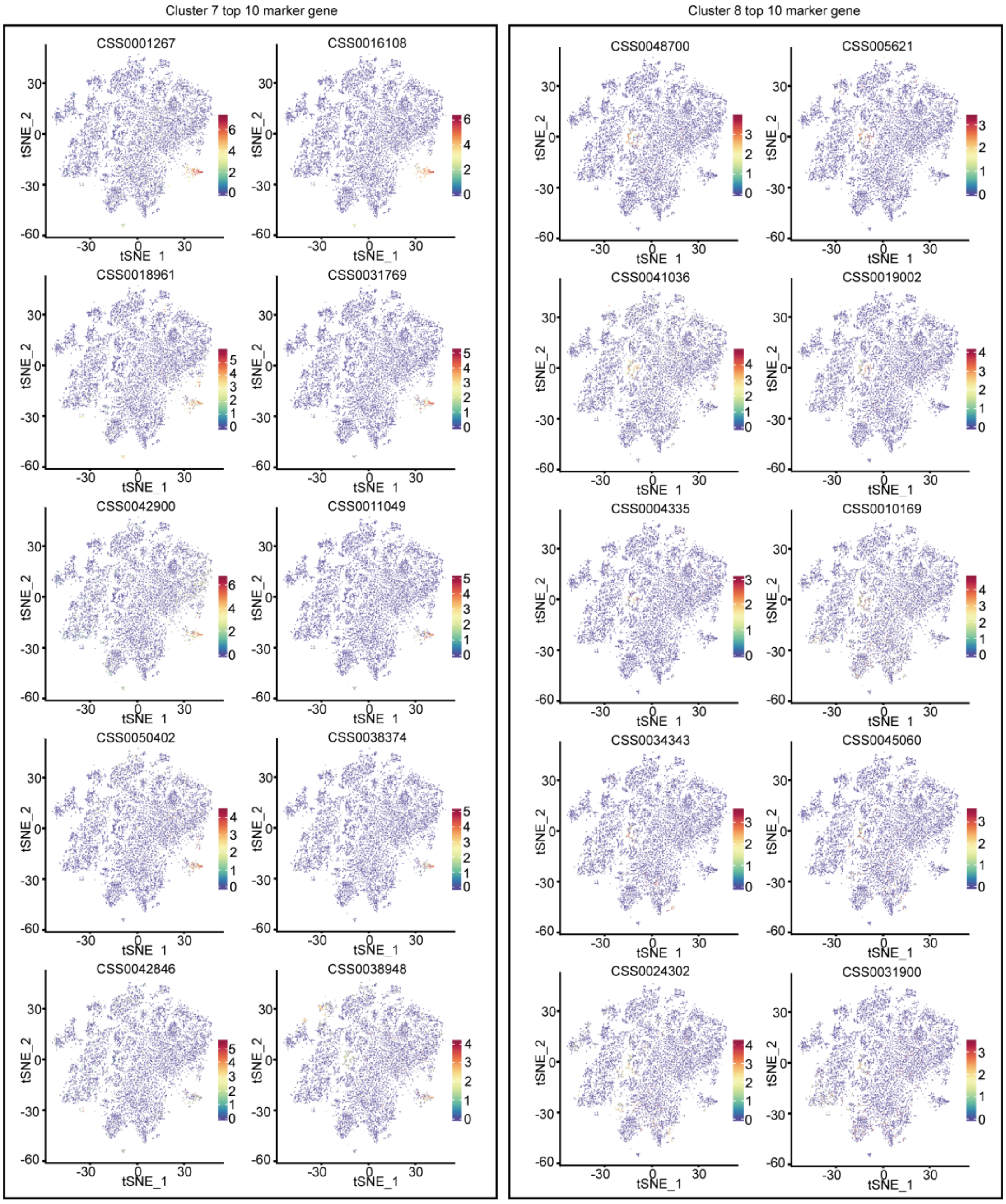


**Supplemental Figure 6. *t*-SNE visualization of cluster 7 and cluster 8 top 10 marker genes.** *t*-SNE plots shows the transcript accumulation of cluster 7 and cluster 8 top 10 marker genes in individual cells. Color intensity indicates the relative transcript level for the indicated gene in each cell.

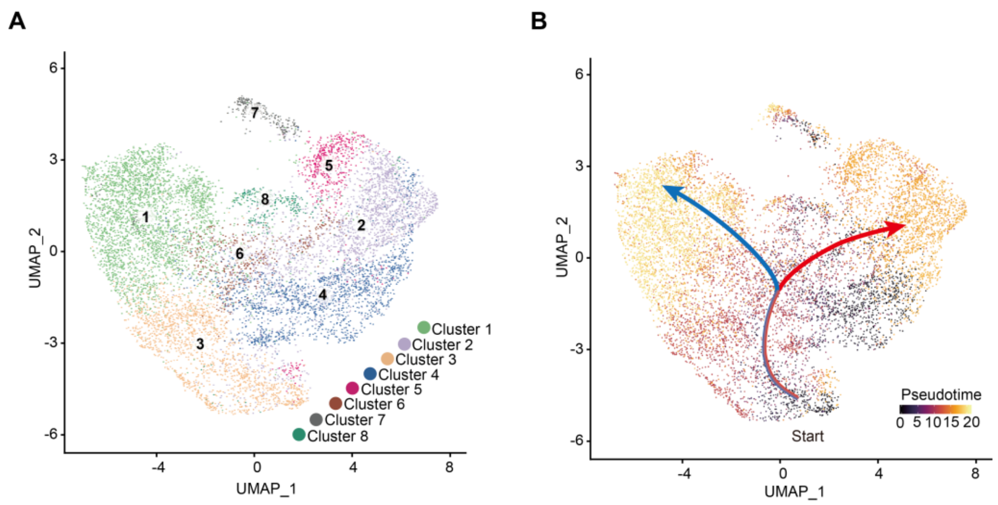


**Supplemental Figure 7. UMAP visualization of eight clusters and differentiation trajectory atop the UMAP. (A)** UMAP visualization plot of 10,435 tea plant root cells. Each dot denoted a single cell. **(B)** Simulation of the successive differentiation trajectory of tea plant root cells over pseudo-time. Colors of the dots represent the pseudo-time score. Red and blue line mark major differentiation trajectories.

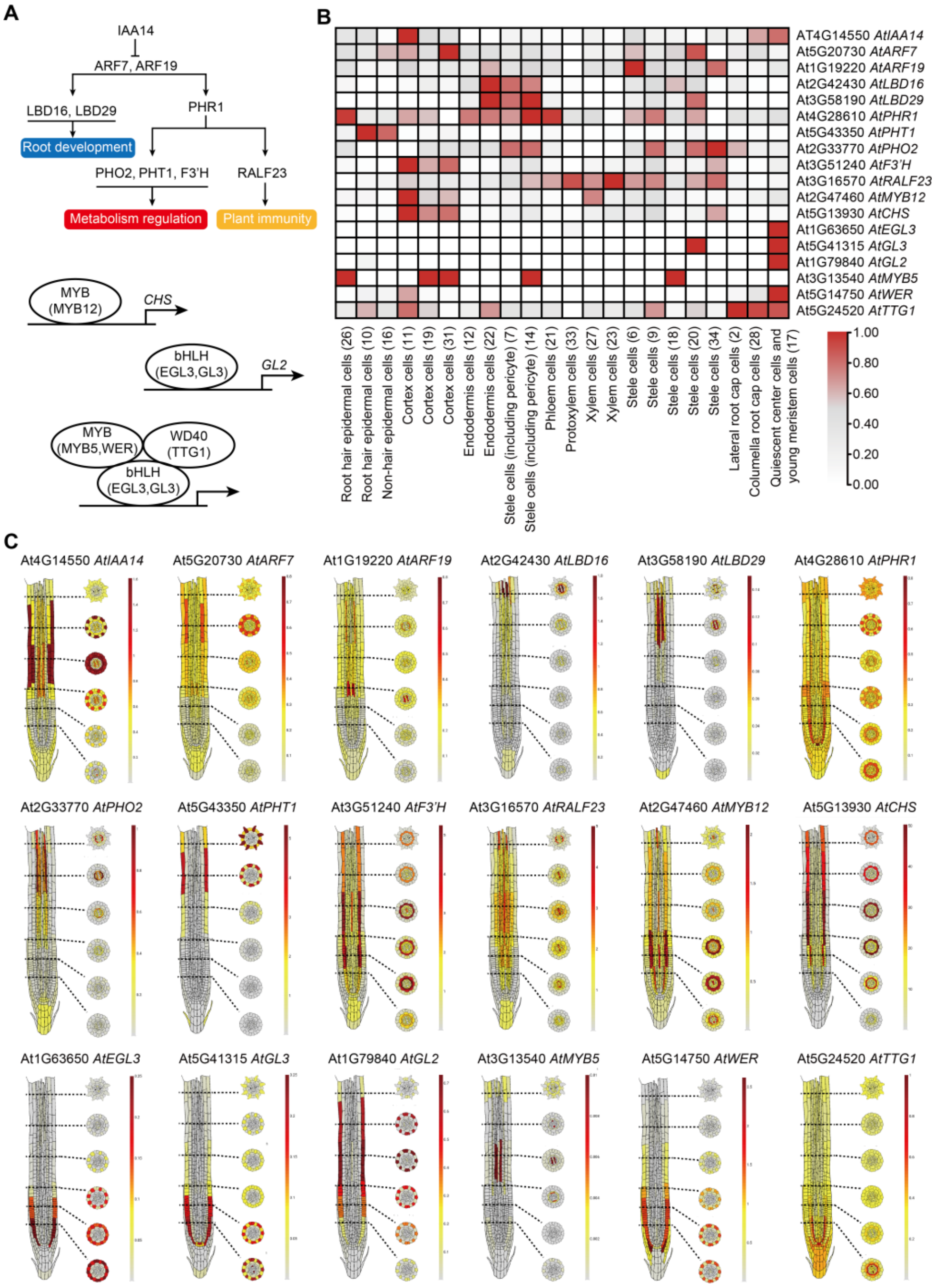


**Supplemental Figure 8. Cell cluster assay of regulators and target genes based on published *Arabidopsis thaliana* data. (A)** Schematic diagram of reported gene regulatory relationships. A hierarchical regulatory network with IAA14 as the upstream regulator (top). MYB-bHLH-WD40 (MBW) complex regulates downstream gene (bottom). **(B) and (C)** Heatmap (B) and tissue-located heatmap (C) for cell cluster expression patterns of genes which were showed in schematic diagram of (A). The scRNA-seq data of *Arabidopsis thaliana* root was obtained from publicly available data (Ryu et al., 2019; Wendrich et al., 2020; Zhang et al., 2019; Denyer et al., 2019; Jean-Baptiste et al. 2019; Shulse et al., 2019; Shahan et al., 2022)

and database (Root Cell Atlas, https://rootcellatlas.org/; BAR, https://bar.utoronto.ca/#GeneExpressionAndProteinTools). The pictures of (C) were generated from Root Cell Atlas web with slight modification.

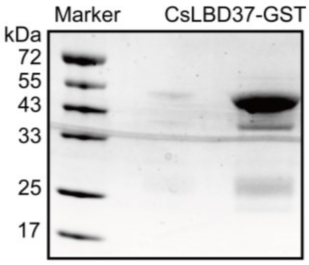


**Supplemental Figure 9. Protein purification of *CsLBD37*.** Protein purification of *CsLBD37*. The CsLBD37-GST fusion protein has a molecular weight of 50.9 kDa (i.e., 24.9 kDa + 26 kDa).

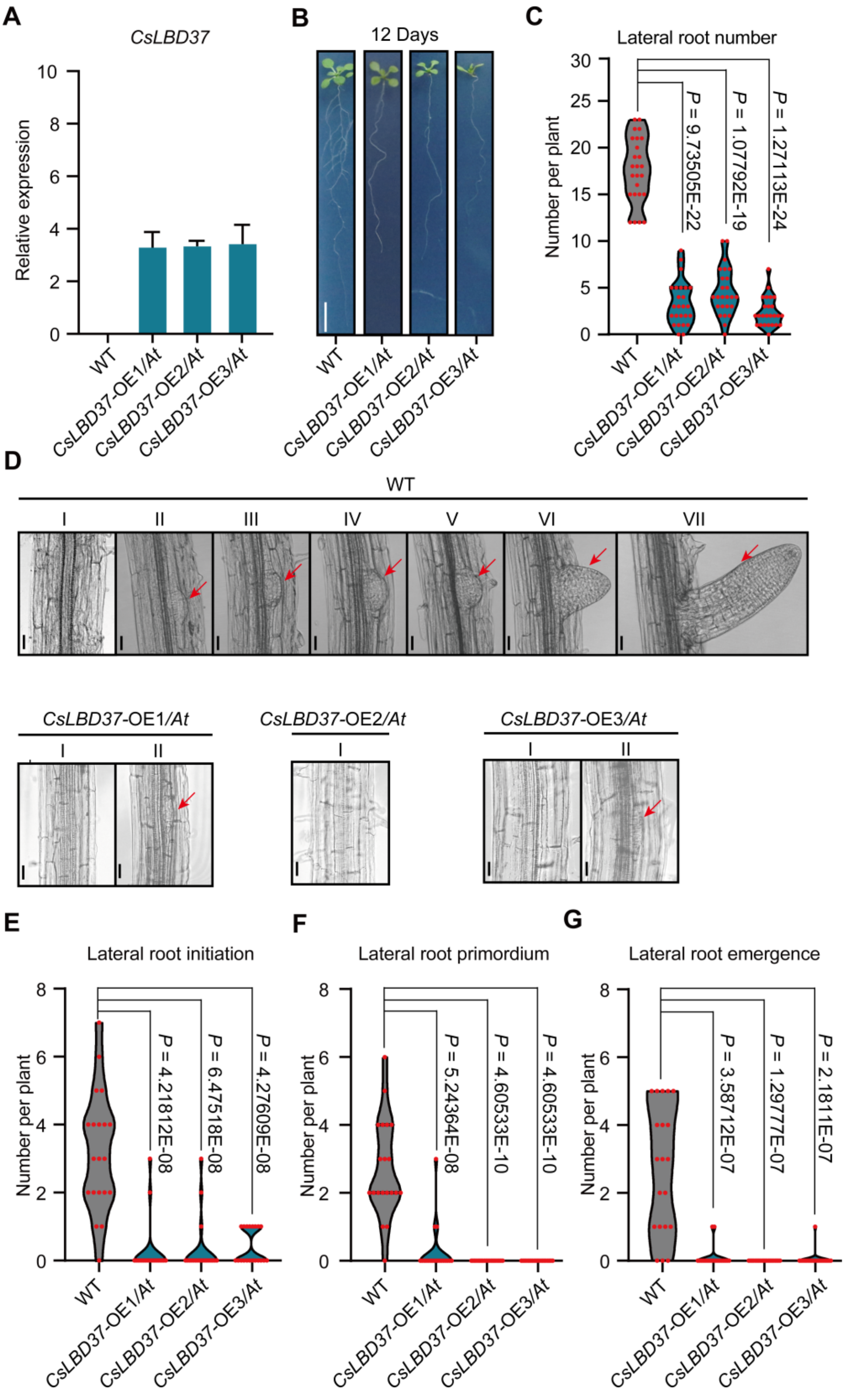


**Supplemental Figure 10. Overexpressing *CsLBD37* in *Arabidopsis* inhibited lateral roots development. (A)** qRT-PCR shows transcription level of *CsLBD37* in three transgenic *Arabidopsis* lines (*CsLBD37*-OE1/*At*, *CsLBD37*-OE2/*At*, *CsLBD37*-OE3/*At*) and wild type *Arabidopsis* (WT). **(B and C)** The growth phenotype of *CsLBD37* transgenic *Arabidopsis* lines and WT in MS medium for 12 days. Scale bar = 1 cm. Lateral root number, each line contained 25 independent biological replicates **(C)**. **(D-G)** The lateral root development phenotype of *CsLBD37* transgenic *Arabidopsis* lines and WT on the 5th day after germination. The phenotype of *CsLBD37* transgenic *Arabidopsis* lines and WT under microscope. Scale bar = 25 μm **(D)**. Lateral root initiation number **(E)**; lateral root primordium number **(F)** and lateral root emergence number **(G)**, each line contained 20 independent biological replicates. **(C and E-G)** Significant difference was determined by two-sided Student's *t*-test.
